# A novel, precise and high-throughput technology for viroid detection in cannabis (MFDetect™)

**DOI:** 10.1101/2023.06.05.543818

**Authors:** Angel Fernandez i Marti, Marcus Parungao, Jonathan Hollin, Berin Selimotic, Graham Farrar, Tristan Seyler, Ajith Anand, Riaz Ahmad

**Affiliations:** Department of Environmental Science, Policy and Management, University of California, Berkeley, Berkeley, CA, United States; MyFloraDNA, Inc. 1451 River Park Dr, 95815 Sacramento, California, United States; Glass House Farms, 645 W Laguna Road, Camarillo, California, United States

**Keywords:** hop latent viroid, cannabis, viroids, RT-LAMP, qRT-PCR, MFDetect™, cannabis pathogen

## Abstract

Hop latent viroid (HLVd) is a severe disease of cannabis, causing substantial economic losses in plant yield and crop value for growers worldwide. The best way to control the disease is early detection to limit the spread of the viroid in grow facilities. This study describes MFDetect™ as a rapid, highly sensitive, and high-throughput tool for detecting HLVd in the early stages of plant development. Furthermore, in the largest research study conducted so far for HLVd detection in cannabis, we compared MFDetect™ with quantitative RT-PCR in a time course experiment using different plant tissue, leaf, petiole, and roots at different plant development stages to demonstrate both technologies are comparable. Our study found leaf tissue is a suitable plant material for HLVd detection, with the viroid titer increasing in the infected leaf tissue with the age of plants. The study showed that other tissue types, including petiole and roots, were equally sensitive to detection via MFDetect™. The assay developed in this research allows the screening of thousands of plants in a week. The assay can be scaled easily to provide growers with a quick turnaround and cost-effective diagnostic tool for screening many plants and tissue types at different stages of development.

## Introduction

Hop latent viroid (HLVd) is an emerging disease in cannabis that significantly affects plant vigor, yield, and bud quality. The viroid was first found and isolated from two commercially grown hop cultivars in North Spain (Pallas et al., 1987). It has been reported that although HLVd-infected hop plants are asymptomatic, it can cause substantial yield loss and decrease both bitter acid contents and terpenes (Faggioli et al., 2017). HLVd has also been found in Stinging Nettle *(Urtica dioica*) and two cultivars of hop, including commercial hop (Humulus lupulus) and Japanese hop (Humulus japonicus Sieb. and Zucc.). The viroid was initially reported in cannabis in 2019 in California from two independent studies (Bektas et al., 2019; Warren et al., 2019). A 2021 survey from Dark Heart Nursery Research concluded that 90% of cannabis facilities in California were infected with HLVd. Their recent report states that HLVd is a hidden threat and most likely the biggest concern for cannabis and hop growers worldwide (https://mjbizdaily.com/experts-sound-alarm-over-global-spread-of-cannabis-viroid/). There are estimates that HLVd could result in crop loss of up to USD 4 billion annually for the cannabis industry (Adkar-Purushothama et al., 2023).

Viroids are the smallest known protein-free infectious RNA molecules that predominantly cause plant disease (Diener, 1971). They rely on host-cell DNA-dependent RNA polymerase and processing enzymes for replication and pathogenesis (Semancik, 2003). HLVd is a small, circular RNA molecule with 256 nucleotides, including a conserved central region (CCR) and terminal conserved hairpin (TCH) (Puchta et al., 1988; Adkar-Purushothama et al., 2023). The symptoms produced by the viroid depend upon the host plant species and viroid variant (Flores et al., 2005). In cannabis, the viroid is associated with “dudding” or “duds” disease of cannabis, which is one of the most devastating cannabis diseases (sy. Hemp) (*Cannabis sativa, Cannabis indica* and *Cannabis ruderalis*). Once infected, the viroid spreads rapidly through *Cannabis* plants, leading to stunted growth, weaker flower smell, diminished flower quality, lower yield, and up to 50% reduction in cannabinoid and terpene production (Adkar-Purushothama et al., 2023, Warren et al., 2019, Singh and Singh 202, Bektas et al., 2019).

HLVd is very stable in plants and is easily transmitted by vegetative propagation or mechanically through contaminated tools (Pethybridge et al., 2008). Due to the non-coding nature of viroids, they recruit host DNA-dependent RNA polymerase II for replication. The resulting replication is error-prone, creating “quasi-species” or sequence variants (Flores et al., 2005). Comparative analysis of all HLVd sequences isolated from cannabis plants identified two distinct HLVd isolates, isolates Can1 (GenBank Acc. No.: MK876285) and Can2 (GenBank Acc. No.: MK876286) (Warren et al. in 2019). The Can1 isolate has 100% sequence similarity with HLVd species, while the Can2 isolate has a point mutation, U225A (Warren et al., 2019). Interestingly, both isolates are infectious and have been reported from HLVd-infected cannabis plants in the USA (Bektaş et al., 2019, Chiginsky et al., 2021).

The viroid can remain latent, and plants remain symptomless for an extended period. As a result, infected plants are accidentally propagated, spreading the viroid and making it hard to eradicate from infected plants and growing areas. Early screening of plants and implementation of integrated pest management (IMPM) seem to be the most efficient means of limiting the spread of the disease (Pethybridge et al., 2008, Postman and Hadidi, 1994, Howell et al., 1997). The early detection of infected plants and remediation is critical to reducing the rate of infectivity, the spread of the viroid, and minimizing losses.

Several technologies have been developed and used to detect viroids in plants. The methods include bioassay, nucleic acid hybridization (Owens and Diener, 1981), return-polyacrylamide gel electrophoresis (Schumacher et al. 1986), RT-PCR (Boonham et al., 2004, Chandelier et al., 2010, Mascia et al., 2010), and reverse transcription loop-mediated isothermal amplification (RT-LAMP; Notomi et al., 2000, Tsutsumi et al., 2010). Of these, RT-PCR and RT-LAMP are considered to be standard procedures. However, both have their advantages and drawbacks. RT-PCR is a very accurate method for pathogen detection; however, this technique is expensive, requiring sophisticated equipment, expert personnel to perform the test, and it lacks high throughput capability. RT-LAMP is gaining a lot of popularity for pathogen detection in plants, animals, and humans. The advantage of RT-LAMP is its capability for high throughput at low cost, and turnaround times can be rapid (Notomi et al., 2000). It provides simple visual colorimetric detection techniques that bypasses the need to use a sophisticated analytical instrument. RT-LAMP uses four to six primers to target and amplify a specific region of the pathogen, and the reaction occurs isothermally in a single tube. RT-LAMP can be designed for detecting multiple viroids using a generic primer or universal set of primers (Bostan et al. 2004; Tseng. 2021).

The virulent nature of this viroid in cannabis and its high rate of transmissibility and spread across most of the gardens in California demand a novel approach that combines the robust and accurate detection of RT-PCR and the low-cost and high throughput of RT-LAMP to allow containment of viroid spread within California and to other states in the country and cultivation areas worldwide. Here, we describe such an approach that MyFloraDNA, Inc. has developed and named MFDetect™. Furthermore, there is currently an open public debate about what is the best plant tissue for pathogen detection so in this study we want to resolve this by testing our approach and qRT-PCR on a range of plant tissues.

Therefore, the objectives of this study include: 1) developing a hybrid method combining the ease of RT-LAMP and sensitivity of RT-PCR for highest accuracy, high-throughput, and low cost, 2) comparing the accuracy of MFDetect™ with qRT-PCR for HLVd detection, 3) identifying a preferred tissue type for early viroid detection during development of the plants.

## 2. Material and Methods

### 2.1. Development of MFDetect™ Method

MFDetect™ is a proprietary technology developed by the MyFloraDNA research team for plant pathogen detection. This innovative testing platform combines RT-LAMP and RT-PCR to facilitate the accurate identification of numerous pathogenic viruses, viroids, and fungi. For HLVD detection, we designed primers (Table 1) using the whole genome sequence for Hop Latent Viroid retrieved from the NCBI database (<u>MK795612.1</u>). The primers were designed (Primer Express 3.0) to work at high annealing temperatures, resulting in high specificity of target amplification while eliminating nonspecific targets. In addition, the RT-PCR validation step enhances detection and quantification of pathogen load in infected plants similar to the CT value associated with qRT-PCR analysis. Primers were synthesized by Integrated DNA technology and prepared in nuclease-free water and stored at –20°C and at a concentration of 16 µM.

**Table 1:**
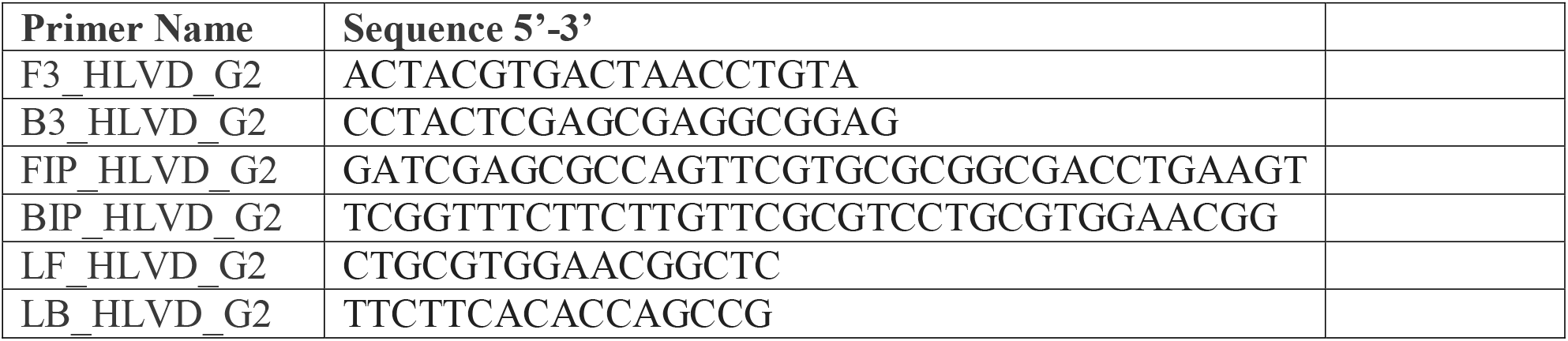
Primers used to amplify HLV in Cannabis plants in this study.

Initially, these primers were tested on synthetic DNA designed from the HLVd genome and subsequently HLVd-positive and HLVd-negative plant samples were confirmed visually by our nursery collaborators. After confirming that the primers only amplified the HLVd genome and there was no nonspecific amplification of other viruses, viroids, or cannabis genome, the primers were selected for testing HLVd in cannabis at large scale.

For MFDetect™ protocol, leaf, petiole, and root tissues were collected in 2ml Eppendorf tubes. Two beads (3.5mm) were added to each tube and the tissue was ground to powder in liquid nitrogen using a TissueLyser (Qiagen) for 5 minutes. Five hundred microliters of unique RNA Extraction Buffer, developed by MyFloraDNA, was added immediately to preserve the RNA integrity. Extracted RNA was diluted further at 1 to 50 ratio using nuclease-free water. From the diluted RNA, 5µl were used for cDNA conversion and amplification of HLVd genome at the 20µl reaction volume using a commercial master mix and enzymes (ThermoFisher), and proprietary primers.

Reactions were prepared in 96 well plates with two HLVd positive controls and two no-template controls (NTC) in each plate. After an initial amplification, PCR products were validated and quantified further by Vii 7 qPCR (Applied Biosystem) using FAM channel as reporter dye and Low Rox as a passive dye to normalize the reporter signal. Delta RN values were calculated to quantify the magnitude of the fluorescence signal generated during the PCR at each time point. Delta RN for MFDetect™ was calculated and compared in three plant tissues i.e., leaf, petiole, and root.

### 2.2. Validation of MFDetect™ by qRT-PCR

For HLVd testing with TaqMan qRT-PCR (Norgen, Canada), we extracted RNA from leaf, petiole and root tissue using RNeasy Plant Mini kit (Qiagen), quantified RNA with NanoDrop and used 15ng of RNA in a total volume of 20µl reaction mixture using the manufacturer’s protocol and run through Vii 7 qPCR (Applied Biosystem). In each 96 well plate, we ran 4 HLVd positive and 2 HLVd negative samples as control. VIC was used as a passive reference dye to normalize the reporting signal of the FAM dye. Ct values were recorded when the positive samples crossed the threshold level.

We collected leaf tissue from 5030 cannabis plants, including multiple cultivars, and at different plant growth stages (5-10 weeks old) from our partner Glass House Farms. After an initial screening of these 5030 plants, we selected forty-four HLVd-positive plants and six HLVd non-infected plants from the 5–6-week-old stage and tested them again using the classic RT-PCR protocol.

### 2.3. Identifying a preferred tissue type for early viroid detection during development of the plants

We aimed to determine the variability of the HLVd load in three different tissue types, leaves, petioles, and root tissue. For that, we collected leaves, petioles and roots and placed them in separate tubes from each of the fifty plants making a total of 150 samples to be tested. In addition to testing different tissue types, we wanted to analyze the same plant material using two different technologies, TaqMan RT-PCR (Norgen, Canada) and MFDetect^TM.^, and for this we duplicated the number of samples to 300 (150 for RT-PCR including roots, petioles, and leaves and 150 for MFDetect™ including roots, petioles and leaves).

To understand the progression and virulence of the pathogen through the growth of the plant, we used the same parameters as in the previous experiment that included the three different plant tissues (leaf, petiole and root) but adding this time a new hypothesis –how soon and in what concentration we can detect the pathogen? For this, the experimental design included three different ages of the plants (5-6 weeks old, 7-8 weeks old and 9-10 weeks old) making a total of three-hundred samples to be analyzed. Considering all the parameters and hypotheses, within this study, a total of 600 samples were analyzed, which represents the largest research study conducted so far for HLVd detection in cannabis.

## 3. Results

### 3.1 Development and Optimization of the MFDetect™ assay

MFDetect™ primers amplified only in the HLVd positive controls as well as in the synthetic HLVd DNA, but not in the HLVd negative samples nor in the cannabis genome. We also optimized the amount of RNA for the RT-LAMP reaction by running a serial dilution experiment and selected the 40 times dilution as the best for RT-LAMP reaction. We tested four different dyes to visualize and quantify the amplification products. We picked one that gave us confirmed positive and negative results that can be validated by RT-PCR as seen in Figure 1.

**Figure 1:**
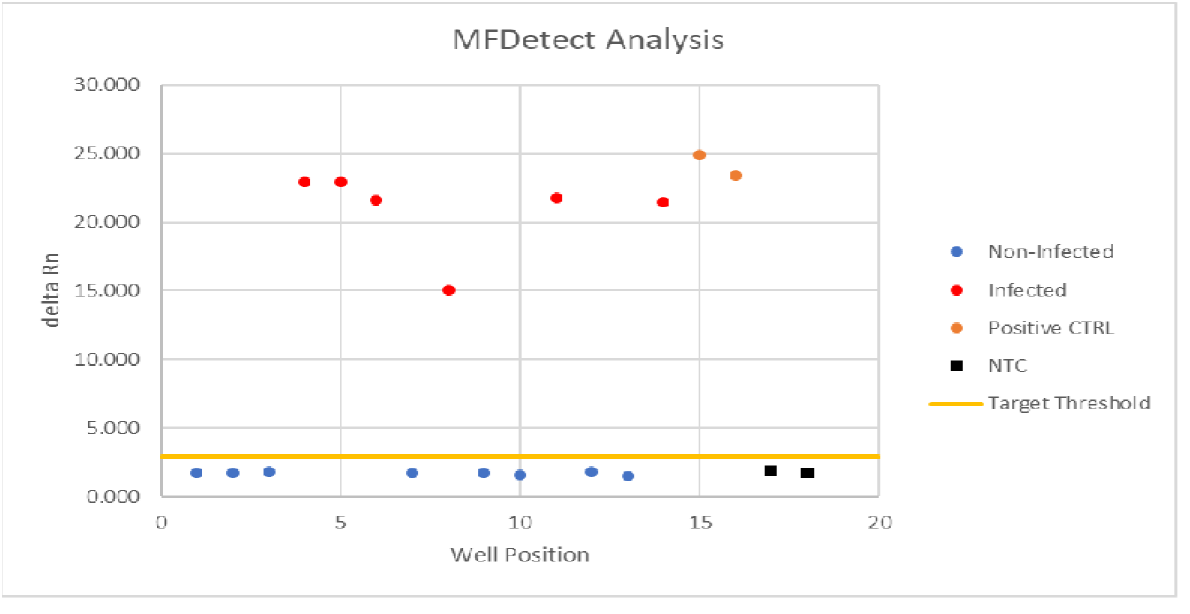
Plot representation of positive and negative detection by using *MFDetect*^*TM*^.

### 3.2 Comparison of MFDetect™ and RT-PCR

To further validate MFDetect™ sensitivity and accuracy in detecting HLVd, we conducted side-by-side experiments comparing MFDetect™ and qRT-PCR techniques in forty-four infected and six non-infected plants. The ΔRn and CT values from a selected number of plants were plotted to create the standard curves as shown in Figure 2. The standard curves for the different samples with MFDetect™ and qRT-PCR were similar. As expected, the infected samples had higher ΔRn values (MFDetect™) or lower Ct values (qRT-PCR), while the uninfected plant samples were below the detection limits. The data demonstrates MFDetect™ is highly accurate and sensitive for detecting HLVd-infected plants. The ease of sample preparation, lower cost, and high throughput make MFDetect™ a powerful HLVd detection technique for viroid presence in plants.

**Figure 2:**
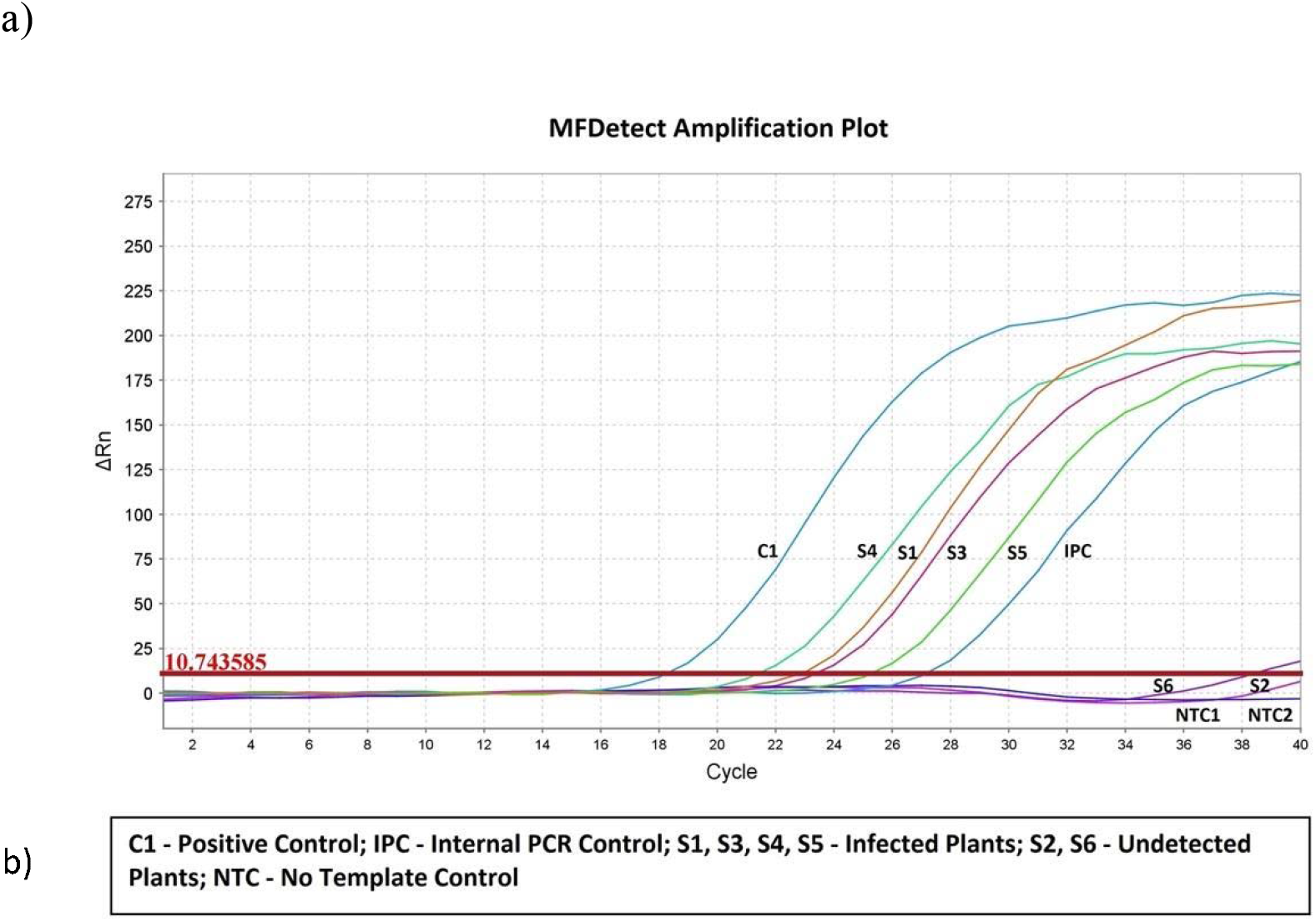
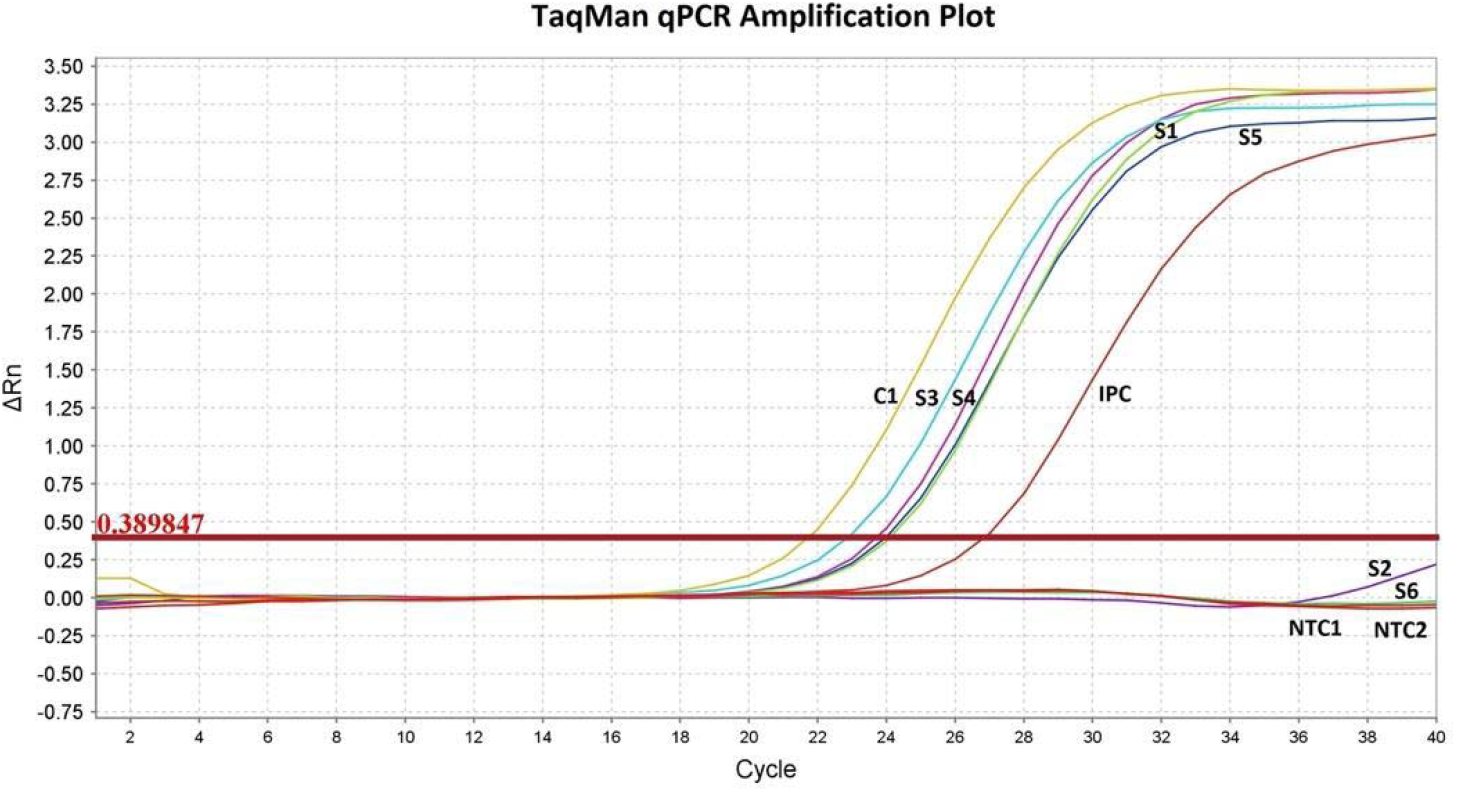
Standard curve for MFDetect™ and RT-PCR using the leaf samples. Amplification plots showing the; a) standard curve of the ΔRn values and b) standard curved of the CT values of the infected (S1, S2, S3, S4), uninfected (S5, S6), positive syn DNA control (C1,), internal positive control (IPC), no-template control (NTC) used in the experiment.

### 3.3 Identifying a preferred tissue type for early viroid detection during plant development

To determine the presence and variability of the viroid by tissue type and age of the plants, three different tissue types-leaf, petiole, and roots-were sampled biweekly from the forty-four infected and six uninfected plants. All the tissue types were collected in duplicate to compare side-by-side MFDetect™ and qRT-PCR. A total of 600 samples were analyzed in this study and the data are summarized in Table 2.

**Table 2.** Comparison of the different plant parts and viroid progression with plant age using MFDetect™ and TaqMan qRT-PCR, with their standard error (SE).

| Tissue Type | Set 1 (5-6 wk) MFDetect™ | Avg $\Delta Rn$ | Set 2 (7-8 wk) MFDetect™ | Avg $\Delta Rn \pm SE$ | Set 2 (7-8 wk) TaqMan RT-PCR | Avg CT $\pm SE$ | Set 3 (9-10 wk) MFDetect™ | Avg $\Delta Rn \pm SE$ | Set 3 (9-10 wk) TaqMan RT-PCR | Avg CT $\pm SE$ |
| --- | --- | --- | --- | --- | --- | --- | --- | --- | --- | --- |
| Leaf | 44 | 12.8 | 44 | 24.2 ± 1.2 | 43 | 21.9 ± 0.35 | 44 | 38.1 ± 2.6 | 44 | 19.7 ± 0.5 |
| Petiole | 44 | na | 43 | 27.1 ± 2.0 | 43 | 21.3 ± 0.5 | 43 | 26.1 ± 2.2 | 43 | 21.8 ± 0.3 |
| Root | 44 | na | 41 | 23.8 ± 1.2 | 44 | 21.5 ± 0.3 | 43 | 19.7 ± 1.6 | 44 | 23.0 ± 0.3 |
| Total | 132 |  | 128 |  | 130 |  | 130 |  | 131 |  |

With MFDetect™ we noticed infectivity in 132/132 (100%) of the leaf samples, 86/88 (98%) of the petioles, and 84/88 (95%) in the root samples, respectively. While for the qRT-PCR assay, the infectivity ranged from 87/88 (99%) in leaves, 86/88 (98%) in petioles, and 88/88 (100%) in roots. The above results indicated that leaves are equally suitable for detecting HLVd from the infected plants compared to petioles and roots. Our results further strengthened the accuracy of MFDetect^TM,^ achieving >99% accuracy compared to qRT-PCR (Table 3).

**Table 3.** Data comparing the accuracy of the MFDetect™ on leaf samples collected from the forty-four infected and six uninfected plants with plant age.

| Plant Type | Tissue | Set 1 (5-6 wk)<br>MFDetect™ | Set 2 (7-8 wk)<br>MFDetect™ | Set 3 (9-10 wk)<br>MFDetect™ | MFDetect™<br>Consistency (Set1,<br>Set2 & Set 3) |
| --- | --- | --- | --- | --- | --- |
| Infected | Leaf | 44 | 44 | 44 | 99.9% |
| Uninfected | Leaf | 6 | 5 | 5 | 83.3% |
| Inconsistent | Leaf | 0 | 1 | 1 | -- |
| Total |  | 50 | 50 | 50 |  |

The viroid load in the infected plants by tissue type and age was determined by the comparing the average ΔRn and Ct values, respectively. We noticed an apparent increase in the ΔRn values for the leaf material from 5-6-week-old material to the 9-10-week plants. The increase in ΔRn values ranged from 1.9X (7–8-week-old plants) to 3X (9–10-week-old plants) compared to values in the samples from the 5-6-weeks old plants. Thus, suggesting the viroid load in the leaf samples accumulated and amplified over time. A similar trend was observed with the CT values in the duplicated leaf samples analyzed with qRT-PCR. A 2.2 average CT value drop in 9–10-week-old plants was observed compared to the CT values in the 7-8-week-old plants. Which again reinforces that the viroid load increased in the leaves over time. However, no such increase in viroid load was observed in the petiole and root tissues (Table 2). The above observation further suggests that the viroid replicates in the leaf tissue and is uploaded into other parts as the plant ages. Our study indicates leaf is a good candidate and most likely the ideal tissue for HLVd detection with the MFDetect™ and qRT-PCR.

Concerning the analysis conducted to investigate the infection rate during the plant development, our results obtained from one hundred fifty leaf samples (132 from infected and 18 from uninfected plants) indicates that the 132 infected samples matched 99.9% when compared over time (5-6, 7-8 and 9-10 weeks old). Whereas sixteen out eighteen (89%) matched for the uninfected plants. Our data indicated that MFDetect™ achieved 99.9% accuracy in detecting infected plants as early as week 5-6 and then corroborated the presence of the viroid at a higher level at week 7-8 and 9-10. In one sample, we found inconsistency. For the analysis conducted at week 5-6, one plant sample in the subsequent analysis at weeks 7-8 and 9-10 weeks was found negative. We speculate the viroid titer in the plant sample was too low, outside the detection limit of the assay.

As we can observe in Figure 3, the HLVd titer in the leaf tissue of the infected plants increased with the age of the plants. To determine the viroid titer in the infected tissue as plants aged, we compared the average ΔRn values for the 5–6-week-old plants with the ΔRn values of the 7-8 week and ΔRn values of the 9-10 week. The ΔRn values increased over time, reaching 1.9X to 3X higher in the older plants. The higher ΔRn values correspond to the higher amounts of the viroid in the older leaf tissue, suggesting the viroid replicated and amplified in the leaves. Based on these results, we inferred that the higher load of the viroid makes the leaf an excellent material for HLVd detection. The presence of the viroid in three different tissue types at different titers (ΔRn and CT values) suggests the viroid load varies with the tissue type and the plant’s age. Our studies further reinforce that once infected, HLVd is persistent in cannabis with the potential to be detected from any plant parts with both techniques.

**Figure 3:**
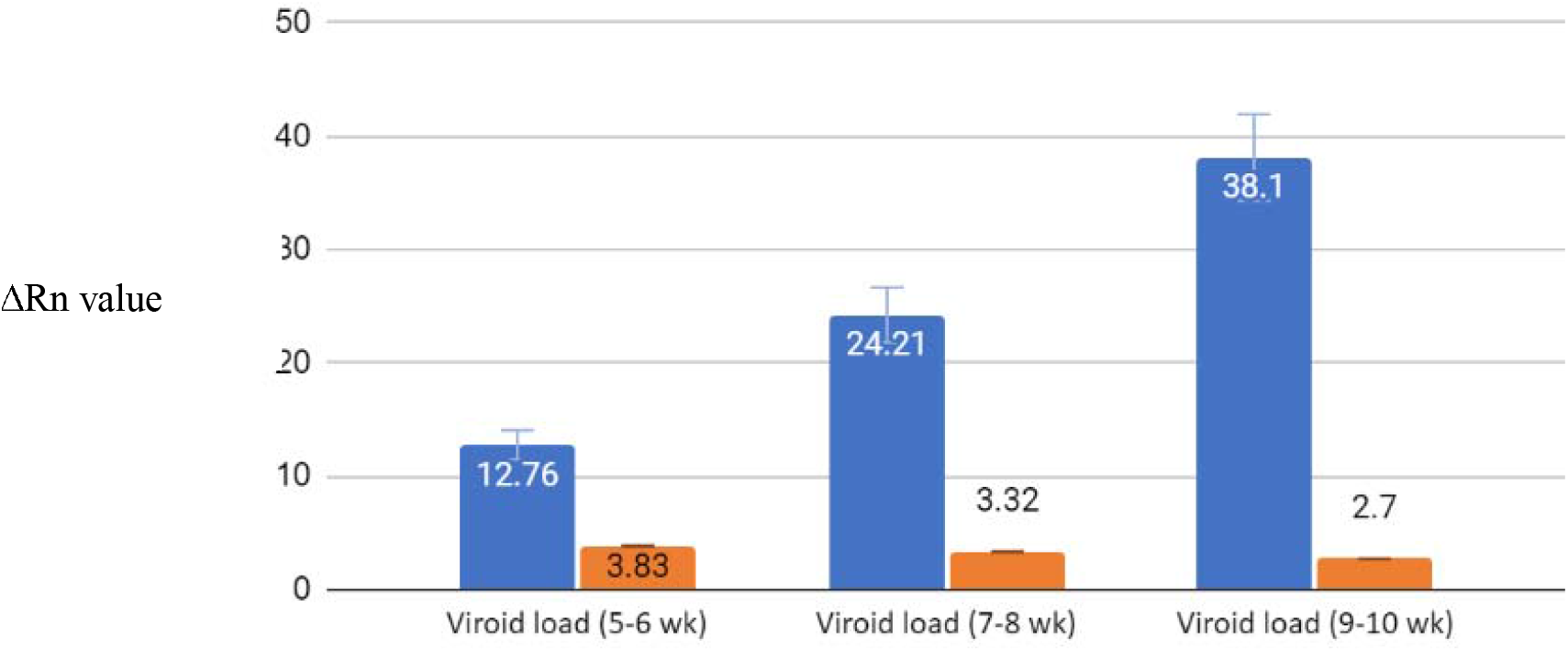
Histogram with the average ΔRn values in the 44 infected (blue bars) and six uninfected (orange bars) plants at different time points starting from 5–6-week-old plants to 7-8-week-old to 9–10-week-old plants.

## 4. Discussion

Hop latent viroid (HLVd) is the biggest threat to cannabis growers. The viroid can spread rapidly via mechanical transmission or by unsterilized farm tools, leading to stunted growth, reduced yields, leaf malformation, terpene damage, diminished flower quality, and susceptibility to other pathogens (Adkar-Purushothama et al., 2023, Massart et al., 2017, Singh, 1970). HLVd-infected plants can be asymptomatic for a long time; its worldwide occurrence raises great concerns for transmission, as do most viroids (Adkar-Purushothama et al., 2023). Regular inspection and healthy cultivation management practices are required by the growers to mitigate the large-scale dissemination of the diseases. This becomes critical based on the report that >90% of the cannabis cultivated in California is infected with HLVd. One of the biggest challenges facing the cannabis industry is the timing and identification of infection to minimize the spread of the viroid (Nachappa et al., 2020, Punja, 2021). Thus, a reliable, high throughput sensitive method at lower cost is needed for large-scale testing of the plants in a grow facility and to restrict the spread of viroids. Additionally, the screening will, in turn, produce healthy and high-quality plants for growers.

Various techniques have been used to detect viroid in infected plants, including RT-PCR and LAMP (Mascia et al., 2010, Notomi et al., 2000, Tsutsumi et al., 2010). The RT-PCR assay has been reported to detect different viroids in different plant species (Nakahara et al., 1999, Luigi and Faggioli, 2011, Luigi and Faggioli, 2013, Malandraki et al., 2015, Hagemann et al., 2021). Although the efficiency of this assay is up to the mark for the detection of viroids (Roslan et al., 2023), the specificity of primers and probes makes it complex and not cost-effective for large-scale detection (Gardner et al., 2003, Nagy et al., 2017). This study describes the MFDetect™ system that relies on a unique set of primers and one-step RT-LAMP plus RT-PCR for detecting HLVd in infected plants. A set of oligonucleotides was designed using the program (Primer Explorer 3.0) to detect HLVd successfully. The optimized primer design and concentration in our RT-LAMP reaction significantly improved the specificity and expanded the capacity to detect and inspect plant material.

Previous reports have shown that the addition of intercalating dyes improves the interpretation of the LAMP data and allows real-time monitoring of the amplification reaction. However, the nonspecific binding of these dyes to end products could also increase the risk of false positive calls (Motoki et al., 2009). Thus, we tested several different dyes in our studies in combination with a new RNA extraction buffer to improve specificity and lower the presence of contaminants to improve RT-LAMP detection. In addition, RNA dilutions up to 40-fold were found to reduce the inhibitory compounds in the crude RNA extracts. The combination of single-step RT-LAMP and calorimetric detection has been widely used for virus detection in plants and for COVID-19 diagnosis (Ali et al., 2023, Alhamid et al., 2023, Bostan et al., 2004; Tangkanchanapas et al., 2018, Tseng et al. 2021). To date, we have analyzed over 150,000 samples using MFDetect™ to detect several other pathogens, including viruses and fungi. We report the most extensive sample used for the detection of multiple pathogens to our knowledge while writing this report.

The sensitivity and accuracy of the MFDetect™ assay were further validated by comparing it with qRT-PCR. Of the fifty plant samples received from our partners, 49 and 50 samples respectively were correctly identified as positive or uninfected with RT-LAMP and qRT-PCR methods, respectively. The discrepancies could be because of the extremely low load of the viroid in one of the uninfected plant samples, which could be beyond the detection limits of the RT-LAMP assay. The sensitivity of the RT-LAMP for the positive samples was comparable to the qRT-PCR. Similar observations comparing the sensitivity of RT-LAMP to RT-PCR have been previously reported (Warghane et al., 2017, Zhao et al., 2015). In another study, RT-LAMP was shown to be more sensitive than RT-PCR for the detection of six different viroids in Solanaceae (Tseng et al., 2021). Thus, our findings here are consistent with the previous studies.

Furthermore, a recent report suggested that RT-LAMP was 100 times more sensitive than RT-PCR in detecting the Indian citrus ringspot virus (ICRSV) (Kokane et al., 2021). The consistency of our results indicates no RNA degradation and no inhibitory compounds in the RNA extraction buffer used for the MFDetect™ assays. Because of the high sensitivity, MFDetect™ may be able to detect infection in the leaves at an early stage of growth before symptoms appear, which is currently the limitation in the industry. Our findings show that a simple RNA extraction method with a highly sensitive RT-LAMP, as previously suggested (Singh et al., 2004, Kankone et., 2021), may further improve the application of MFDetect™ for viroid detection in cannabis and hemp.

HLVd is mainly transmitted through infected propagules or mechanically through vegetative propagation, infected tools, and grafting (Barbosa et al., 2005). Few studies have reported viroid transmission through pollen and seed (Mahafee et al., 2009, Pethybridge et el., 2006). After establishing the sensitivity of MFDetect™, we focused our study on the tissue type and age of the plants on determining the best tissue and earliest plant growth stage for HLVD detection. In the study with 600 samples comparing the age and tissue type, we detected the viroid in the leaf samples collected from 5-6 week-old cuttings from mother plants. Our study reports the earliest stage of plant growth for HLVd detection from cannabis. Our future investigation is to enable detection of the viroid in plant cuttings, preferably 1-2 week-old rooted plants. In a subsequent experiment, we observed that leaf, petiole, and root tissue types are equally sensitive to HLVd detection with MFDetect™ and Taqman RT-PCR. The ease of collecting tissue material and the higher viroid loads in the leaf material make it an ideal material for HLVd detection. It is also reported that HLVd is unevenly distributed in the plants. Based on our findings, we suggest sampling leaves from different plant parts to ensure the infected plants do not go undetected. Viroid replicates and moves through phloem trafficking to lower parts of the plants, from source to sink. When the viroid accumulates in the roots, the plants are probably well-grown, too old, and fully infected. The damage may already be caused, and controlling the spread of the viroid is unlikely.

In conclusion, we developed a new hybrid RT-LAMP/RT-PCR detection system, MFDetect^TM,^ for robust detection of HLVd that is rapid, reliable, sensitive, and cost-effective. The assays were validated in one of the most extensive public experiments collected from one of the largest and more reputable grower’s facilities in the USA. Compared to RT-PCR, MFDetect™ was equally sensitive in detecting HLVd from cannabis plants. To address the growing concerns around tissue type for HLVd detection, we found that all three plant tissues (leaf, petiole, and root) showed the presence of the viroid and gave consistent detection. Because of the ease of sampling, accessibility, and minimal chance of contamination, we recommend that leaf tissues are ideal for HLVd detection. The leaf tissue accumulated higher loads of viroid over time based on the ΔRn values. This further strengthens that leaf is an adequate tissue for early detection. We believe that MFDetect™ is a reliable assay for mitigating the dissemination of the viroid. When combined with healthy cultivation practices and meristem culture one can produce viroid-fee planting material for nurseries and growers.

## Acknowledgments

Authors would like to thank Prof. Richard Dodd, Dr. Peter Roberts, Rubina Ahsan, Oscar Illan and Prof. Umar Rao for technical advice and revision of the manuscript.

## Author contributions

Designed the research: AFM, AA, RA. Project execution: MP, JH, BS, RA. Collection of plant material and phenotyping: TS. Wrote the first draft of the manuscript: AFM, AA, RA. All authors reviewed and corrected the final text.

